# Single Channel Strongly-Coupled Geometry Surface Coil for Small Mammal Whole Brain fMRI

**DOI:** 10.64898/2026.08.03.742081

**Authors:** Kyle A. Johnson, Hanbing Lu, Jason W. Sidabras

## Abstract

Single-channel surface coils remain central to rodent MRI, but conventional circular loop designs face an inherent trade-off between surface and depth sensitivity, limiting whole-brain coverage for applications such as resting-state BOLD fMRI. This work introduces a single-channel strongly-coupled geometry surface coil. It consists of a stop-sign shaped loop inductively overcoupled to a nested, three-turn elongated racetrack spiral designed to improve depth sensitivity and thru-plane coverage while remaining robust to variable sample loading. Benchtop characterization across three phantoms of differing size showed the parallel resonant mode and loaded quality factor changed negligibly with loading. In phantom imaging at 9.4 T, the SCG coil achieved in-plane SNR and temporal SNR comparable to, and at shallow depths exceeding, a commercial Bruker 2×2 receive-only rat brain array, while showing substantially more consistent tSNR across loading conditions. The SCG coil also demonstrated superior thru-plane tSNR over a 20 mm slice range at 3.5 mm depth, approximating the anterior-posterior extent of the rat brain. In vivo resting-state BOLD fMRI in eight rats, acquired with a double asymmetric spin-echo EPI sequence, yielded a default mode network consistent with prior reports and revealed a previously undescribed subcortical network spanning superior/inferior colliculi and cerebellar regions. These results establish the single-channel SCG as a promising foundation for next-generation rodent receive coil arrays, combining loading-independent tuning with extended sensitive coverage suitable for whole-brain functional imaging.

## 2 Introduction

Introduced to the MRI community in the 1980s, surface coils and receive coil arrays offer several immediate advantages over their single channel receive coil predecessor, including more complete coverage of the target anatomy and an enhancement in signal-to-noise ratio (SNR) (Ackerman et al., 1980; Kneeland et al., 1986; Roemer et al., 1990). It was later recognized that the spatially varying sensitivities of individual receive coil channels could also be leveraged to accelerate acquisitions (Deshmane et al., 2012). Research to maximize these benefits continues today, with receive coil channel counts around a fixed volume increasing to an unprecedented 128 at both 7 T and 10.5 T (Gruber et al., 2023; Lagore et al., 2025). Paired with the latest image acceleration strategies and image reconstruction algorithms, images are being collected faster and with better spatial resolution and SNR than ever before.

Substantial progress in receive coil array technology has, however, been largely confined to human imaging. The rat brain is orders of magnitude smaller than the human brain, severely limiting how many receive coil channels can be placed around it with commercially-available hardware currently capped at 8 receive coil channels. Scaling channel count down further to fit more around the rat brain is not feasible since 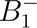 is inversely proportional to the radius of the receive coil channel (Hoult and Richards, 1976; Hoult, 2000). Rather than emphasizing channel count, maximizing sensitivity within each individual channel becomes the priority when building arrays under this constraint.

Currently, single channel receive coils in rodent brain imaging are dominated by circular loops, and while ubiquitously used, they carry well-known limitations. Chief among these is the 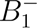 versus loop radius trade-off mentioned earlier (Hoult and Richards, 1976; Hoult, 2000). In practice, smaller loops yield relatively high SNR near the surface of the imaged object but fall with depth, and are often too small to cover the entire rat brain (Gao et al., 2019). Larger loops do recover some SNR at depth and provide full brain coverage, but sacrifice surface SNR compared to their smaller counterparts (Gao et al., 2019). This trade-off has meaningful consequences in imaging applications, such as resting-state blood oxygenation level-dependent (BOLD) functional magnetic resonance imaging (fMRI), when adequate SNR is typically required throughout the rat brain.

Several approaches have been explored to address these limitations. Stretchable receive coils were shown to increase both surface SNR and SNR at depth, but their frequency response and thus their SNR varied with sample geometry (Vishnu Ramesh et al., 2023). Implantable receive coils have also been used and similarly improve SNR at both the surface and depth by placing the coil in closer proximity to the brain, but they require invasive surgery to affix the coil to the skull (Martin et al., 2013; Pirttimaki et al., 2013).

Concentric planar loops, or more generally inductively coupled coil pairs, represent a compelling non-invasive alternative to both stretchable and to implantable receive coil designs in rodent imaging. Early implementations used one loop coupled to a second resonant loop as a means of matching to the 50Ω of the coaxial cable (Mispelter et al., 2006). Other implementations later included dual-tuned planar concentric loops for X-nuclei/^1^H imaging (Takahashi et al., 2023) and concentric planar coil arrays for parallel imaging (Ohliger et al., 2005). Despite this, relatively little work has explored the unique properties that emerge when two resonant loops are overcoupled into what we term the strongly-coupled regime (Sidabras et al., 2016; Terman, 1947). Our group first introduced the strongly-coupled geometry (SCG) in the context of MRI in 2016 (Sidabras et al., 2016), and another group has since independently demonstrated the 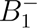 enhancement and thermal noise reduction it yields in a phantom (Li et al., 2025).

Building on our 2016 manuscript, we show here that a stop sign geometry (primary) strongly coupled to an elongated racetrack spiral geometry (secondary) is negligibly affected by loading with different samples and achieves comparable in-plane temporal signal-to-noise ratio (tSNR) and SNR as a function of depth to a commercially-available Bruker ^1^H receive-only 2×2 rat brain surface coil array. We further demonstrate superior thru-plane tSNR over 20 mm at a depth of 3.5 mm in a single phantom, and resting-state BOLD fMRI connectivity maps in 8 rats that reveal distinct brain networks spanning nearly 20 mm, roughly the full length of the rat brain excluding the olfactory bulb. Taken together, these results suggest the SCG could represent the foundation of a next-generation single-channel surface receive coil, with future work aimed at building it into an array that fully unlocks the advantages array technology offers.

## 3 Methods

### 3.1 SCG Construction

The single channel SCG surface coil was built closely following the original 2016 publication (Sidabras et al., 2016). First, the 15×20 mm stop sign (primary) and the 12×18 mm, ∼3 turn, elongated racetrack spiral geometry (secondary) were tuned to *ω_p_* and to *ω_s_*, respectively. Once tuned, the secondary was nested inside the primary. Such close proximity causes the primary and the secondary to strongly couple through mutual inductance (Sidabras et al., 2016; Terman, 1947). Because the system is in this strongly coupled regime, a frequency (mode) split occurs, which is illustrated in Figure 1c. Next, the parallel mode (*ω_∥_*) was tuned to the Larmor frequency via a tuning capacitor connected in parallel with the primary. Then, the strongly-coupled geometry was matched to 50 Ω using a capacitive ladder network. Pin diodes are included for active decoupling. Both the capacitive ladder and the active decoupling circuit topologies are detailed in the 2016 publication (Sidabras et al., 2016). A low-noise amplifier (WanTcom, WMA9RA, Chanhassen, MN, USA) was then connected to the output of the matching network followed by two in-house built baluns placed along the coaxial cable at electrical lengths of *λ/*4 and 3*λ/*4 (Peterson et al., 2003). The fabricated single channel SCG surface coil is shown in Figure 2.

**Figure 1:**
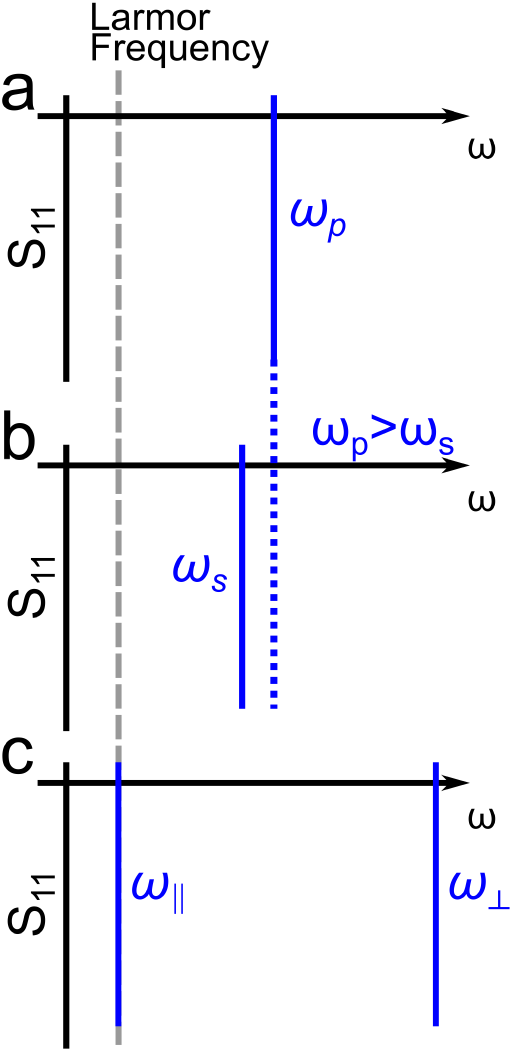
Illustrative representation of tuning the inductively coupled pair. The plots have frequency (*ω*) on the *x*-axis and *S*_11_ on the *y*-axis. Larmor frequency is indicated by the gray dotted line. First, the primary is tuned to *ω_p_* (a). Then, the secondary is tuned to *ω_s_* (b). *ω_s_* is set to be as close as possible to *ω_p_* while *ω_p_ > ω_s_* holds true. Finally, the primary and the secondary are inductively coupled into the strong coupling regime causing the frequency split (c). The parallel mode *ω_∥_* is fine tuned to the Larmor frequency by increasing *ω_p_*via a tuning capacitor.

**Figure 2:**
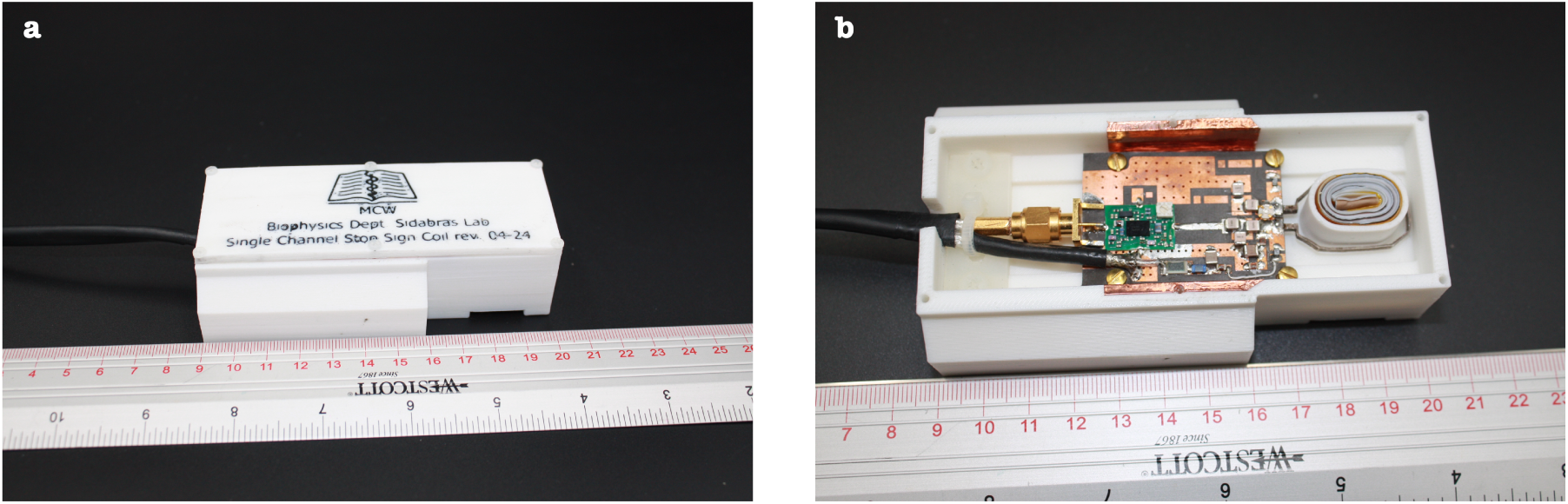
Fabricated single channel SCG surface coil. (a) Resonator housing 3D-printed on a Bambu Lab X1 Carbon with white and green TPU 95A HF filament. The housing was designed to interface with the Bruker animal cradle. (b) Resonator housing without the top cover. Inside is a three-turn, 12×18 mm elongated racetrack spiral strongly coupled to a 15×20 mm stop sign loop. The stop sign loop is soldered to a printed circuit board (PCB) crafted from Rogers 5880. The PCB is affixed to the housing with four set screws. On-board is the capacitive ladder matching network, WanTcom WMA9RA LNA, and active decoupling.

### 3.2 Phantom Fabrication

Three phantom housings were constructed from stock G10 fiberglass tubing (1.250′′ inner diameter, 1.352′′ outer diameter), each cut to a length of 3.089′′. Two of the three housings were further reduced in outer diameter using a lathe, yielding three housings with varying outer diameters: “no trim” (1.352′′), “medium trim” (1.320′′), and “most trim” (1.292′′). Each end of the phantom housings was sealed with a rubber O-ring and an acrylic cap. A hole was tapped through one cap and sealed from the outside with a nylon screw and a rubber O-ring, allowing the phantom to be filled and resealed. A 1 L stock solution was prepared gravimetrically by dissolving 1 g of CuSO_4_ · 5 H_2_O and 3.6 g of NaCl in deionized water to achieve target concentrations of 4 mM and 62 mM, respectively, following a formulation provided by Bruker. The phantom housings were filled from this stock solution via syringe through the tapped hole.

### 3.3 Benchtop Characterization

The single channel SCG surface coil was characterized on the bench by measuring resonant frequency (f_0_) and quality factor (Q) under different loading conditions (unloaded, “no trim”, “medium trim”, and “most trim”) on a vector network analyzer (PNA-X N5224B, Keysight, Santa Rosa, CA, USA). Prior to any data collection, a two port electronic calibration was performed from 300 to 500 MHz at 1 MHz increments (Electronic Calibration Module N4693-60001, Keysight, Santa Rosa, CA, USA). Once calibrated, the single channel SCG surface coil was taped to the top of the “no trim” phantom, 10 VDC was provided to the on-board LNA, and its output was connected to port 2. A 15 mm sniffer loop was connected to port 1, was positioned above the single-channel SCG surface coil such that lowering the sniffer loop any more resulted in a broadening of the S_21_ curve, and that height was fixed for the unloaded, “medium trim”, and “most trim” phantoms. The S_21_ measurement in dB at each frequency was recorded for every loading condition and was outputted as separate comma-separated value (csv) files. The csv files were loaded into Python and S_21_ was plotted as a function of frequency and loading condition. *f*_0_ and Q were defined as arg max[*S*_21_(*f*)] and 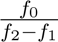, respectively, where *f*_2_ and *f*_1_ are the upper and lower −3dB frequencies.

### 3.4 Phantom Imaging

Phantom imaging experiments were performed on a Bruker BioSpec 94/20 USR 9.4T magnet with AVANCE III-HD hardware and ParaVision 6.0.1 software. A Bruker ^1^H transmit-receive 86 mm volume coil (model: T12054V3) was used for excitation. Signal reception involved one of two receive-only surface coils in separate experiments: 1) Bruker ^1^H receive-only 2×2 rat brain surface coil array (model: T10324) and 2) an in-house fabricated receive-only single-channel SCG consisting of a three-turn elongated spiral geometry nested inside a 15×20 mm stop sign loop. Each receive coil was evaluated with all three phantoms, resulting in six coil–phantom combinations in total. For each experiment, the receive-only surface coil was centered on the phantom and positioned at isocenter. A 2D Fast Low-Angle Shot (FLASH) pulse sequence was run for each coil-phantom combination. Imaging parameters were identical across all experiments and were as follows: 45×45 mm^2^ field-of-view, 256×256 acquisition matrix, 5.1 ms echo time, 400 ms repetition time, 30*^◦^* flip angle, 11 slices, 0.350 mm slice thickness, 3.5 mm slice gap, 55 repetitions, and no Partial Fourier or parallel imaging acceleration.

### 3.5 Signal-to-Noise Ratios

Images were reconstructed in the ParaVision 6.0.1 reconstruction environment. Temporal signal-to-noise ratio (tSNR) was computed voxelwise as the mean of the timeseries signal divided by its standard deviation. Separately, the mean of the timeseries signal per voxel was also divided by the mean temporal standard deviation of voxels within a region-of-interest outside the phantom to yield a voxelwise signal-to-noise ratio (SNR). Both tSNR and SNR as a function of depth per phantom-coil combination were plotted in the middle axial slice along the superior-inferior axis coincident with the center of the vector reception field of the surface receive-only coil. tSNR was also plotted as a function of slice position for each coil-phantom combination at a constant depth of 3.5 mm.

### 3.6 Animal Imaging

All procedures were approved by the National Institute on Drug Abuse (NIDA) Animal Care and Use Committee. Eight adult rats weighing 350–450 g were imaged following an anesthesia protocol described previously (Brynildsen et al., 2017), which has been shown to preserve neurovascular coupling (Fukuda et al., 2013; Lu et al., 2016) and large-scale brain networks (Lu et al., 2012). Rats were induced with 2.5% isoflurane in oxygen enriched air (70% N_2_ + 30% O_2_) followed by a bolus, intraperitoneal dexmedetomidine injection (0.015 mg/kg). Rats were then fastened to a MRI-compatible holder with ear and bite bars, positioned into a nose cone, and a temperature-controlled water heating pad was placed underneath the animal to maintain appropriate body temperature. Once affixed, the isoflurane concentration delivered through the nose cone was gradually lowered and maintained at 0.5%-0.75%. Both isoflurane and expired CO_2_ were actively scavenged through a building vacuum line with the inlet and outlet gases carefully balanced to ensure at least 95% SpO_2_. Dexmedetomidine (0.015 mg/kg/hr) was infused simultaneously with the low-dose isoflurane via a continuous, subcutaneous injection connected to an infusion pump (PHD 2000, Harvard Apparatus, South Natick, MA, USA). Heart rate, respiratory rate, and body temperature were actively monitored (Small animal Monitoring and Gating System, SA Instruments Inc., NY) to ensure stable physiology while imaging.

Imaging experiments were conducted on a Bruker 9.4T magnet equipped with AVANCE-III hardware and ParaVision 7.0 software. A Bruker ^1^H quadrature volume coil (model: MT0381) was used for excitation and the single channel SCG surface coil described above was used for signal reception. For each rat, the imaging study began with a three-plane localizer and a Rapid Acqusition with Relaxation Enhancement (RARE) acquisition. The decussation of the anterior commissure (∼0.36 mm from bregma) appears dark in T2-weighted images. This readily identifiable structure served as the fiducial landmark to standardize intra- and inter-animal slice positioning. Acquisition parameters for the RARE experiment were: 35×35 mm^2^ field-of-view, 256×256 acquisition matrix, 36 ms echo time, 3100 ms repetition time, 8 RARE factor, 31 slices, 1 mm slice thickness, and 0.3 mm slice gap.

After the RARE acquisition and no sooner than 60 minutes following anesthesia induction (Brynildsen et al., 2017), resting-state blood-oxygen level dependent (BOLD) functional magnetic resonance imaging (fMRI) data was collected using a double asymmetric spin echo echo planar imaging (dASE-EPI) pulse sequence recently published by our group (Johnson et al., 2026). Small temporal offsets (*τ*_1_ and *τ*_2_) are introduced to shift the center of the EPI readouts relative to the center of the spin echo created by appropriately spaced 90*^◦^* excitation and 180*^◦^* refocusing radiofrequency pulses. Inclusion of temporal offsets modulates both BOLD contrast and susceptibility-induced signal dropout with the latter being particularly problematic near air-tissue interfaces in the rat brain at ultra-high field. dASE-EPI was shown to mitigate susceptibility-induced signal dropout and maintain reasonable BOLD sensitivity, which prompts its inclusion as a viable resting-state BOLD fMRI acquisition strategy (Johnson et al., 2026). Imaging parameters for the dASE-EPI experiment included: 35×35 mm^2^ field-of-view, 64×64 acquisition matrix, 30 ms spin echo time, 2.18 ms *τ*_1_, 12.33 ms *τ*_2_, 27.82 ms *TE_ASE_*_1_, 42.33 ms *TE_ASE_*_2_, 21 slices, 1 mm slice thickness, 0.3 mm slice gap, 300 kHz receiver bandwidth, and 350 repetitions. Two separate gradient-recalled echo echo planar imaging (GRE-EPI) acquisitions with identical parameters to dASE-EPI, but with opposite phase-encoding gradient polarities, were collected to enable retrospective geometric distortion correction.

### 3.7 Functional Connectivity

Resting state BOLD fMRI data was analyzed using group independent component analysis (ICA) followed by dual-regression to derive statistical functional connectivity maps. EPI images were first subject to identical pre-processing steps reported recently (Johnson et al., 2026). Specifically, geometric distortions in the EPI images were first corrected using data acquired with forward and reverse k-space trajectories using the topup() program in FSL (Smith et al., 2004). The two ASE images obtained within each TR were then averaged to enhance SNR. This averaging was justified because the time between the two ASE images was only 14-15 ms. Given the sluggish nature of the hemodynamic response, any MR signal changes due to spontaneous neuronal activity would be indistinguishable between them. Likewise, motion artifacts arising from cardiac pulsation and respiration would also be effectively identical as their periods are much longer than the time between the two acquisitions. The EPI time series from each rat were then decomposed into spatial independent components using the MELODIC package in FSL (Smith et al., 2004). Noise components that exhibited periodic temporal patterns resulting from respiration and cardiac pulsation were identified and removed. EPI images from the rat that had minimal head tilt and consistent orientation was chosen as the template, to which the EPI images from the other rats were linearly co-registered. The co-registered EPI images from all rats were subsequently averaged to generate a group template specific to this study and the EPI images from each rat were further co-registered to the group template. EPI time series from all rats underwent group ICA using the “concatenation” option with 20 independent components in MELODIC because it produced consistent spatial maps.

Inter-subject analysis and group statistical comparison was performed using the dual-regression approach (Beckmann et al., 2009) as reported previously (Lu et al., 2012). First, individual spatial component maps derived from the group ICA were regressed against each volume of resting-state BOLD fMRI data and the resulting spatial regression coefficients were concatenated across time. Second, the variance-normalized time series from the above spatial regressions were regressed against the corresponding resting-state BOLD fMRI time course to estimate the regression weights of specific component maps in individual animals. Results from all 8 animals were then subject to a one-sample t-test against zero. A threshold of t > 5.0 with the corrected *p <* 0.05 (uncorrected *p <* 0.0015) was applied to generate final maps for each group ICA component. Finally, component maps were co-registered to high-resolution RARE images to aid in structural identification.

## 4 Results

### 4.1 Phantoms Negligibly Effect ***ω_∥_*** and Loaded Q

Without a phantom, *ω_∥_* is 404 MHz and the loaded Q without sample (*Q_wo_*) is 101 as shown in Figure 3a. When loaded with a phantom, both *ω_∥_* and Q decrease (Figure 3b. Specifically, *ω_∥_* is 398 MHz when the single channel SCG surface coil is loaded with the “no trim” phantom. *ω_∥_* decreases further to 397 MHz when the “medium trim” and the “most trim” phantoms load the single channel SCG surface coil. Like *ω_∥_*, the loaded Q with sample of the single channel SCG surface coil is similar between the phantoms. With the “no trim” phantom, loaded Q (*Q_no_*) is 66.3 while the “medium trim” and the “most trim” phantom yield a loaded Q, *Q_med_* and *Q_most_* respectively, of 66.2.

**Figure 3:**
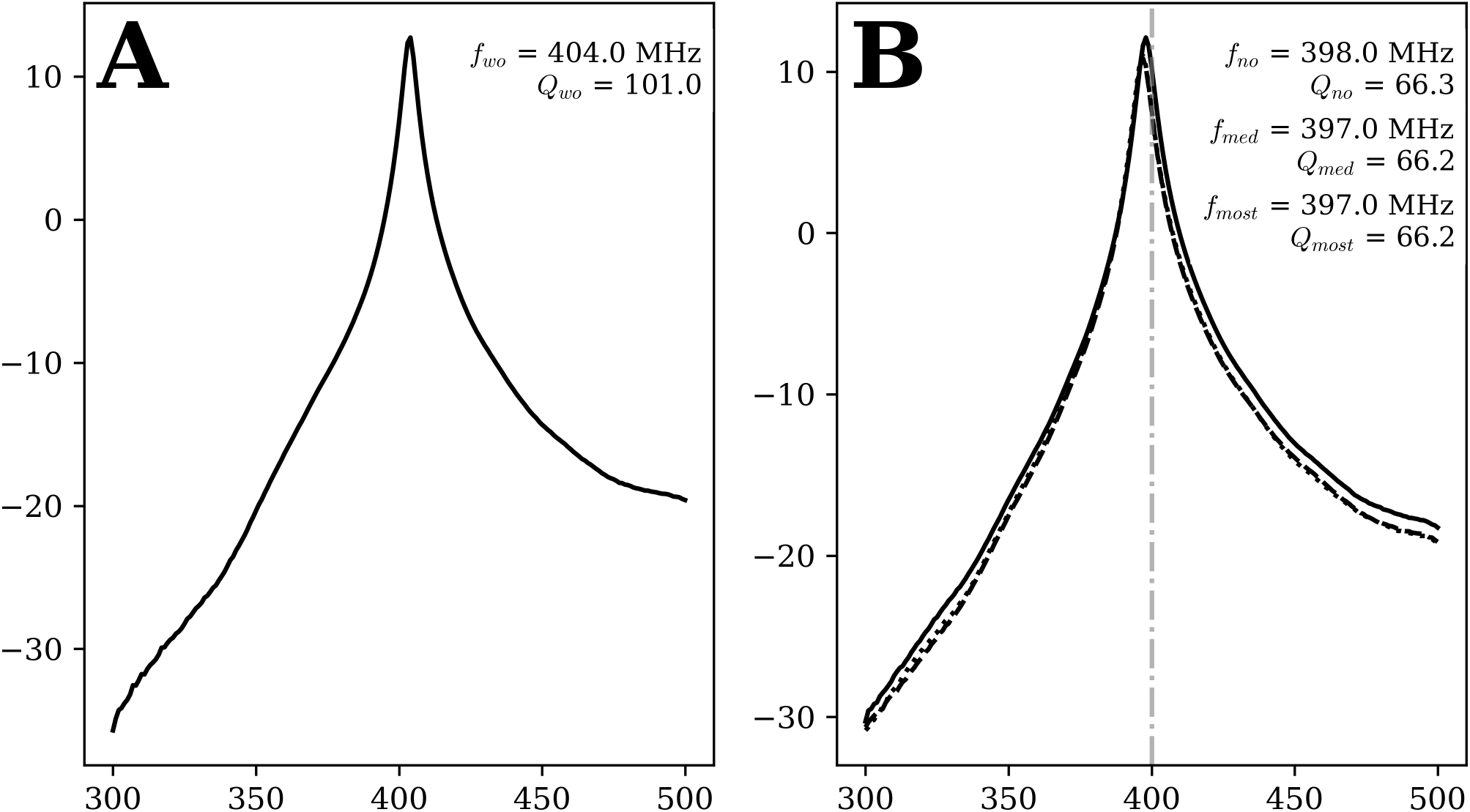
Single channel SCG surface coil S_21_ plots without sample loading (a) and with sample loading (b). Frequency (MHz) is on the x-axis and S_21_ (dB) is on the y-axis. Frequency is swept from 300 to 500 MHz at 1 MHz steps. The gray dashdot line in (b) indicates the Larmor frequency for ^1^H at 9.4T, 400 MHz. Solid, dashed, and dotted lines denote S_21_ responses of the single-channel SCG surface coil with “no trim,” “medium trim,” and “most trim” phantoms, respectively.

### 4.2 Comparable Normalized In-Plane SNR and tSNR

Normalized in-plane SNR is comparable between the single channel SCG surface coil and the Bruker ^1^H receive-only 2×2 rat brain surface coil array. The normalized in-plane SNR of the single channel SCG surface coil, plotted in Figure 4a-c, is ∼20% higher than the Bruker ^1^H receive-only 2×2 rat brain surface coil array at depths up to 4 mm; this holds for each phantom imaged. Beyond 4 mm, the single channel SCG surface coil performs marginally worse than the Bruker ^1^H receive-only 2×2 rat brain surface coil array.

**Figure 4:**
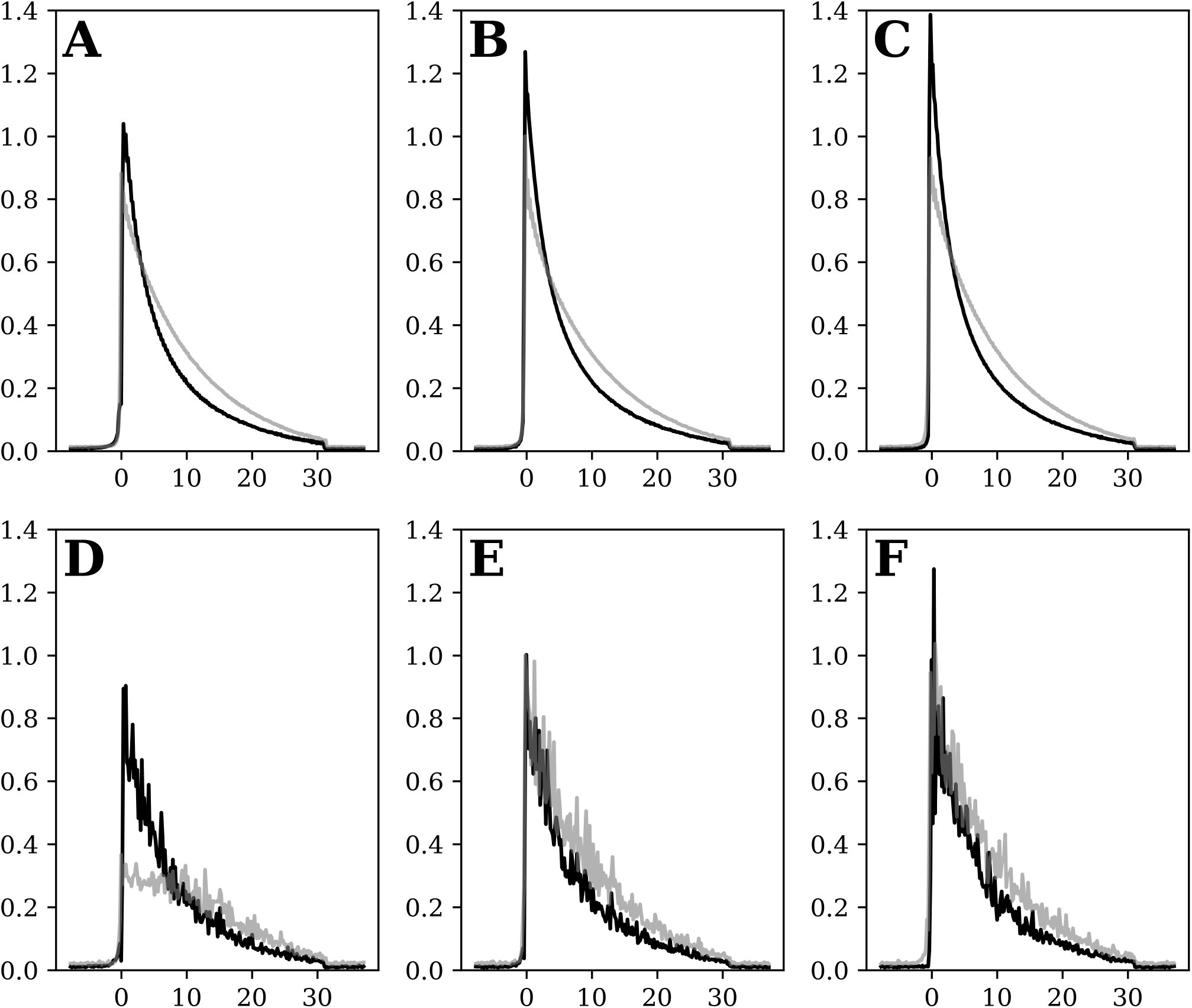
Normalized SNR and tSNR as a function of depth evaluated in the middle axial slice. The x-axis represents depth in mm with 0 mm corresponding to the superior surface of the phantom. Signal is calculated as the mean of the voxel timeseries at depth *d*. The black trace corresponds to the single channel SCG surface coil and the gray trace corresponds to the Bruker ^1^H receive-only 2×2 rat brain surface coil array. In subplots (a)-(c) noise is defined as the mean temporal standard deviation of voxels within a region-of-interest outside the phantom whereas in subplots (d)-(f) noise is defined as the temporal standard deviation of the voxel timeseries at depth *d*. Within each suplot set ((a)-(c) and (d)-(f)), SNR curves are independently normalized to the SNR/tSNR at *d* = 0 from the Bruker ^1^H receive-only 2×2 rat brain surface coil array loaded by the “medium trim” phantom.

In terms of normalized in-plane tSNR, the single channel SCG surface coil performs similarly across the three phantoms as shown in Figure 4d-f. Unlike the single channel SCG surface coil, the Bruker ^1^H receive-only 2×2 rat brain surface coil array has notable normalized in-plane tSNR fluctuations depending on the phantom imaged.

In particular, normalized in-plane tSNR is ∼70% lower when imaging the “no trim” phantom (Figure 4d) compared to the “medium trim” (Figure 4e) and the “most trim” (Figure 4f) phantoms. When comparing normalized in-plane tSNR between the single channel SCG surface coil and the Bruker ^1^H receive-only 2×2 rat brain surface coil array, the single channel SCG surface coil has a higher normalized in-plane tSNR at depths up to 15 mm.

### 4.3 Enhanced Thru-Plane tSNR Over 20 mm in the “No Trim” Phantom

Thru-plane tSNR was also compared between the single channel SCG surface coil and the Bruker ^1^H receive-only 2×2 rat brain surface coil array. When loaded with the “no trim” phantom and at a depth of 3.5 mm, the single channel SCG surface coil has a ≥ 2× thru-plane tSNR over a slice coverage of 10 mm compared to the Bruker ^1^H receive-only 2×2 rat brain surface coil array as shown in Figure 5. Furthermore, the single channel SCG surface coil has higher thru-plane tSNR than the Bruker ^1^H receive-only 2×2 rat brain surface coil array over a slice coverage of 20 mm.

**Figure 5:**
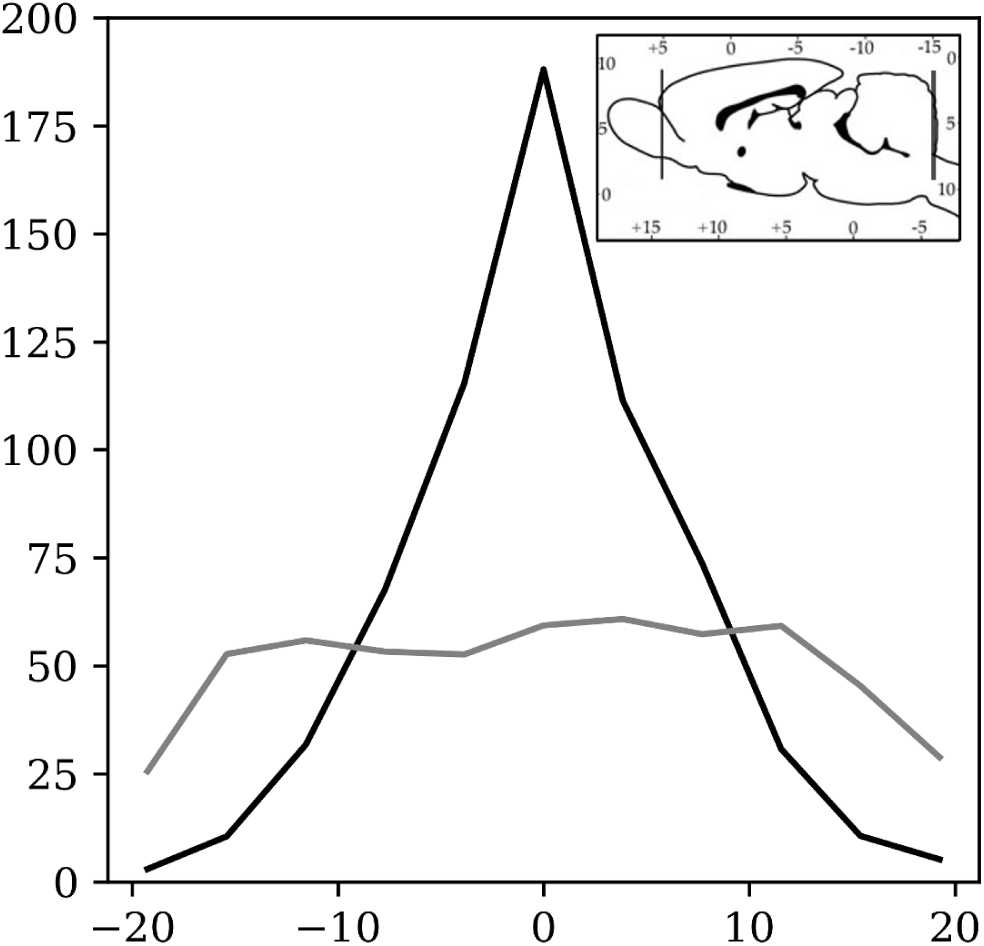
tSNR as a function of slice position at a depth of 3.5 mm relative to the superior surface of the “no trim” phantom. The x-axis is slice location in mm and the y-axis is tSNR. Slice location is defined in mm relative to the center axial slice (slice thickness = 0.35 mm, slice gap = 3.5 mm) with the center axial slice coincident with the center of the vector reception field of the surface coils. The thru-plane tSNR for the single channel SCG surface coil is shown in black and the thru-plane tSNR for the Bruker ^1^H receive-only 2×2 rat brain surface coil array is shown in gray. Superimposed onto the tSNR plot is an illustrative, sagittal slice of a rat brain with a mm scale.

### 4.4 Large Scale Resting-State Brain Networks

We next performed in-vivo experiments to examine the spatial coverage and sensitivity of the RF coil. Imaging data were acquired using the double asymmetric spin echo EPI sequence (Johnson et al., 2026). We previously showed that this sequence allowed for partial recovery of MRI signal in regions with severe *B*_0_ field inhomogeneity, such as the cerebellum and amygdala. Figure 6 shows the default mode network (DMN). Significant clusters include the orbital cortex (Orb), prelimbic cortex (PrL) and medial Orb (MO), cingulate cortex (CG1, CG2), retrosplenial cortex (RSG), temporal association cortex (TeA) and secondary visual cortex (V2L). These regions are broadly consistent with our previous report (Lu et al., 2012).

**Figure 6:**
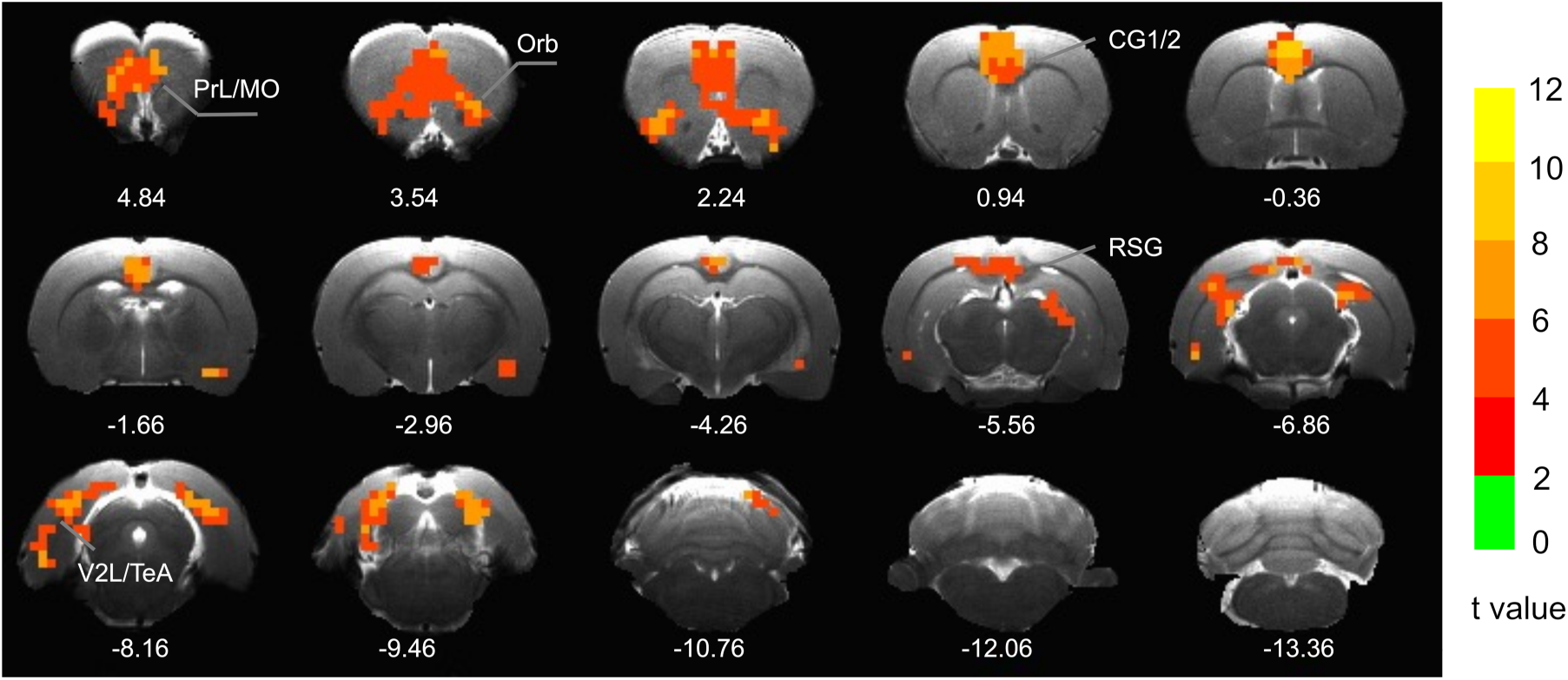
The default mode network in the rat brain identified using group ICA followed by dual regression. Significant clusters include the orbital cortex (Orb), prelimbic cortex (PrL) and medial Orb (MO), cingulate cortex (CG1, CG2), retrosplenial cortex (RSG), temporal association cortex (TeA) and secondary visual cortex (V2L). Color bar indicates t-statistics corrected for multiple comparisons.

In addition to the above DMN, we also identified a subcortical network that incorporates the posterior part of the rat brain. Results are shown in Figure 7. Significant clusters include the superior gray, optical nerve, and intermediate gray layer of the superior colliculus (SuG, OP and InG), dorsal and external central nucleus of the inferior colliculus (DCIC, ECIC), anterior cerebellar lobule (3, 4,5) and cerebellar lobule 6 (Paxinos and Watson, 1998).

**Figure 7:**
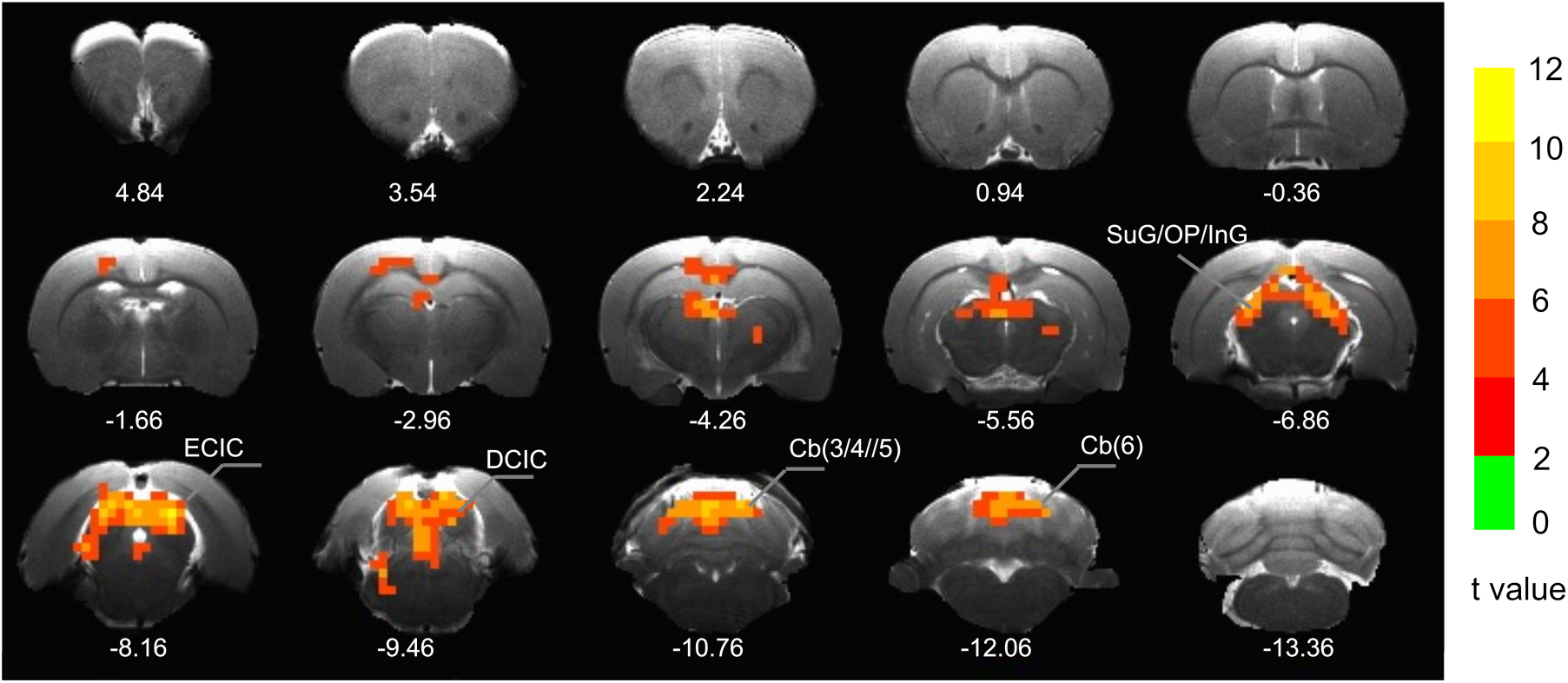
A subcortical network that incorporates the posterior part of the rat brain. Significant clusters include the superior gray, optical nerve, and intermediate gray layer of the superior colliculus (SuG, OP and InG), dorsal and external central nucleus of the inferior colliculus (DCIC, ECIC), anterior cerebellar lobule (3, 4,5) and cerebellar lobule 6. Color bar indicates t-statistics corrected for multiple comparisons.

## 5 Discussion

Across the phantoms, *ω_∥_* and loaded Q remained nearly identical. This consistency is evident in Figure 3b and is likely attributable to mutual inductance. When the primary (stop sign) and the secondary (elongated spiral racetrack) interact through mutual inductance, the impedance of the secondary is coupled onto the primary and appears as an additional series impedance (Sidabras et al., 2016; Terman, 1947). Therefore, when the SCG is excited at resonance with a real frequency *ω*_0_, the input impedance at the primary capacitor then becomes proportional to *Q_L_* and is defined in Eq 1.

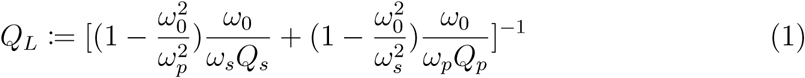

Notably *Q_L_* is a function of *ω*_0_, Q of the primary (*Q_p_*) and Q of the secondary (*Q_s_*), which has implications for when the SCG is loaded with a sample. In the current configuration, the primary sits ∼7 mm above the sample whereas the secondary is situated ∼2 mm above the sample, the thickness of the case. Because of this configuration, it is conceivable that when a sample is introduced the secondary is reasonably loaded while the primary remains relatively unperturbed, owed to their relative distances away from the sample.

On the bench, with the primary positioned ∼7 mm above the sample, *Q_p_* remains reasonably constant across the phantoms, measured as a 5.9% change from the “no trim” to the “most trim” phantom reported in Supplemental Table 1. Comparatively, the secondary at ∼2 mm above the sample has a 21.2% change in *Q_s_* from the “no trim” to the “most trim” phantom. Because *Q_L_* is a non-linear function of both *Q_p_* and *Q_s_*, the theoretical percent change in *Q_L_* (13.2%) is smaller than the change in *Q_s_* because *Q_p_* is relatively constant. This suggests that the mutual inductance inherent to the SCG makes it less susceptible to sample loading. For example, in order for the SCG to be dominantly loaded by the sample with a fixed *Q_p_*, the secondary has to be overcoupled to the sample. A contour plot theoretically demonstrating the relationship between *Q_p_*, *Q_s_*, and *Q_L_* is provided in Supplemental Figure 1.

Relative insensitivity to different loading conditions could be quite beneficial for longitudinal imaging. In both human and animal imaging, variable tuning and matching capabilities are typically excluded. It is assumed tuning and matching on the bench with a phantom is an adequate representation of how samples will load surface coils in the magnet. This assumption is bolstered by the inclusion of arrays having a sufficient number of channels that overcome any SNR penalty associated with an individual element not being precisely tuned and matched. While largely true in human imaging, animal imaging is not as fortunate because arrays tend to have far fewer channels. It is plausible that the number of channels does not overcome the SNR penalty caused by elements not being sufficiently tuned and matched, which could be further exacerbated by day-to-day changes in loading.

This is evident in the normalized tSNR plots in Figure 4(d-f). The Bruker ^1^H receive-only 2×2 rat brain surface coil array exhibits a ∼70% difference in normalized in-plane tSNR between the “no trim” and the “medium trim” phantoms while normalized in-plane tSNR is similar between the “medium trim” and the “most trim” phantom. Unlike the Bruker ^1^H receive-only 2×2 rat brain surface coil array, the single channel SCG surface coil is more robust to different sample loading conditions. Variability in normalized in-plane tSNR can be quite detrimental in longitudinal studies, particularly in experiments where consistent tSNR is important, like BOLD fMRI. Although researchers do their best to build animal cohorts that have minimal age, weight, genetic, and physiological differences, it is not always possible and can become even more diverse once pathology is introduced. Not to mention further complications arise when imaging longitudinally. All of these can possibly contribute to variable loading conditions and a surface coil less susceptible to this then becomes quite valuable.

Complimentary to the consistent, normalized in-plane tSNR is the single channel SCG surface coil having a greater thru-plane tSNR than the Bruker ^1^H receive-only 2×2 rat brain surface coil array, shown in Figure 5, over 20 mm at a depth of 3.5 mm in the “no trim” phantom. This finding is partly driven by the Bruker ^1^H receive-only 2×2 rat brain surface coil array having reduced sensitivity in this phantom, but importantly the geometries of the primary and of the secondary also play a substantial role. The primary is a 15×20 mm stop sign geometry with ∼18 mm of the side rungs having 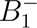 perpendicular to the main magnetic field *B*_0_ compared to the original publication where the primary was a 25 mm circular loop with 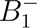 having a cosine dependence relative to *B*_0_ (Sidabras et al., 2016). Furthermore, the secondary is a ∼3 turn, 18 mm long spiral racetrack geometry that also contributes a sensitivity enhancement that is proportional to the number of turns multiplied by the current ratio 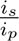 (Sidabras et al., 2016). The geometry of the primary and of the secondary individually, along with the inductive coupling between them, yields a surface coil that has sufficient sensitivity to cover ∼20 mm, which is roughly the length of the rat brain excluding the olfactory bulb and is illustrated in Figure 5.

The in-plane sensitivity (Figure 4) and the thru-plane coverage (Figure 5) together enable the single channel SCG surface coil to be used in whole brain resting-state BOLD fMRI experiments. The DMN depicted in Figure 6 is broadly consistent with the original report (Lu et al., 2012). However, the original DMN map also included the dorsal hippocampus, a region that did not reach statistical significance in the present study. This discrepancy is likely attributable to differences in sample size and data volume. Specifically, the original DMN was identified using 16 rats with a total of 119 echo-planar imaging (EPI) scans, whereas the current study employed 8 rats with only 8 scans.

The subcortical network illustrated in Figure 7 represents a novel finding. The intermediate gray layer of the superior colliculus receives convergent inputs from the cortex and basal ganglia and plays a key role in mediating motor behaviors, including orienting responses such as head and eye movements. The inferior colliculus serves as a higher-order auditory processing center, receiving substantial descending projections from the auditory cortex. It integrates complex auditory and non-auditory (multi-modal) inputs, including somatosensory signals, and contributes to auditory–motor integration. The anterior cerebellar lobules (3,4,5) are primarily involved in sensori-motor integration and motor coordination, while cerebellar lobule VI is critical for fine sensorimotor control and visually guided orientation. Collectively, the network in Figure 7 suggest a functional system supporting multimodal sensory processing and sensorimotor coordination.

## 6 Conclusion

We have demonstrated that a single-channel SCG surface coil, built from a stop-sign loop strongly coupled to an elongated racetrack spiral, offers a compelling alternative to conventional single-loop and commercial array coils for rat brain imaging. The mutual inductance between the primary and secondary elements renders the resonant frequency of the coil and loaded Q largely insensitive to sample variation, a property that translated into markedly more consistent normalized tSNR across differently sized phantoms than was observed with a commercial Bruker 2×2 receive-only array. This loading insensitivity is particularly valuable for longitudinal and cross-subject studies, where day-to-day and animal-to-animal variability can otherwise degrade data consistency. In addition to this robustness, the coil achieved comparable or superior in-plane SNR and tSNR and superior thru-plane tSNR over roughly 20 mm, matching the anterior-posterior extent of the rat brain excluding the olfactory bulb. In vivo resting-state BOLD fMRI using this coil recovered a default mode network consistent with previously published work, along with a novel subcortical network spanning the superior and inferior colliculi and cerebellar lobules, underscoring the sensitivity of the coil across both cortical and deeper brain structures. Together, these findings support the SCG as a strong candidate building block for next-generation single-channel and multi-channel rodent receive coils, where maximizing per-channel sensitivity, rather than increasing channel count, is the more tractable path to improving whole-brain rat MRI/fMRI. Future work will focus on extending this single-channel design into a multi-element SCG array to realize the additional benefits of parallel imaging and further gains in SNR.

## Notes

### Competing Interest Statement

The authors have declared no competing interest.

### Funding

- National Institute of Allergy and Infectious Diseases: 75N93023C00014

